# Enhanced stimulus-induced and stimulus-free gamma in open-eye meditators

**DOI:** 10.1101/2024.02.19.581028

**Authors:** Ankan Biswas, Srishty Aggarwal, Kanishka Sharma, Supratim Ray

**Affiliations:** IISc Mathematics Initiative, Department of Mathematics, Indian Institute of Science; Bangalore, 560012, India; Department of Physics, Indian Institute of Science; Bangalore, 560012, India; Centre for Neuroscience, Indian Institute of Science; Bangalore, 560012, India

**Keywords:** Meditation, Gamma, 1/f slope, EEG

## Abstract

Visual stimuli induce “narrowband” gamma oscillations (30-70 Hz) that are linked to attention/binding and attenuate with aging and neurodegeneration. In contrast, meditation increases power in a broad frequency range (>25 Hz). However, the effect of meditation on stimulus-induced gamma is unknown. We recorded EEG from meditators and controls performing open-eye meditation while gamma-inducing stimuli were presented before, during and after meditation. We found that stimulus-induced gamma, like stimulus-free gamma, was stronger in meditators. Interestingly, both gamma signatures co-existed during meditation but were unrelated and prominent in occipital and fronto-temporal regions, respectively. Further, power spectral density (PSD) slope, which becomes shallower with aging, was steeper for meditators. Meditation could boost inhibitory mechanisms leading to stronger gamma and steeper PSDs, potentially providing protection against aging and neurodegeneration.

**One line summary:** Stimulus-induced and stimulus-free gamma are stronger in open-eye meditators.

## Introduction

Presentation of certain visual stimuli such as bars (*1*), gratings (*2*, *3*) or reddish hues (*4*, *5*) induces a “narrowband” gamma rhythm in the visual cortex, which has a distinct bump in the power spectral density (PSD) of brain signals with a bandwidth of ∼20 Hz and center frequency between 30-70 Hz. This rhythm is modulated by attention (*6*, *7*) and memory (*8*), linked to processes such as binding (*1*, *9*), gain control (*10*) and normalization (*11*), and thought to reflect interactions between excitatory neurons and parvalbumin positive (PV+) interneurons (*12*, *13*). Recent studies have shown that presentation of large (full-screen) stimuli induce a distinct, slower gamma between 20-35 Hz (termed slow gamma), which remains coherent over larger distances compared to the traditional (fast) gamma (*3*), and could reflect the involvement of the somatostatin inhibitory network (*14–16*). Slow gamma is in the same frequency range as beta (12-30 Hz), but it is distinct from the classical beta rhythm which is induced in the sensory-motor areas, prominent during spontaneous period and suppressed by motor movements (*17*). Instead, slow gamma is weak or absent during spontaneous period, induced by visual stimuli and has similar properties and spatial location as fast gamma (*3*). Both slow and fast gamma weaken with age (*18*) and are abnormal in patients suffering from brain disorders like Alzheimer’s Disease (AD) (*19*) and Schizophrenia (*20*). In addition, connectivity across brain regions reduces with onset of AD predominantly in slow gamma (*21*).

Interestingly, gamma power is shown to be elevated in long-term meditators (*22–24*). However, this “stimulus-free” endogenous gamma has a distinct spectral signature. It is characterised by an increase in power over a large frequency range above ∼25 Hz, but there is no distinct “bump” in the PSD (*24*). Importantly, no study to our knowledge has studied the effect of meditation on the stimulus-induced gamma oscillations. Further, since meditation is often performed with closed eyes, it is unknown whether stimulus-induced narrowband and stimulus-free broadband gamma can co-exist, and if so, whether they interact. To address these questions, we recorded electroencephalography (EEG) signals from long-term (>5 years) practitioners of the Brahmakumaris (BK) Rajyoga meditation which is performed with open eyes, and their age and gender matched controls, while gamma-inducing stimuli were shown before, after and during meditation.

## Results

Subjects meditated with open eyes twice – once without any stimulus (protocol M1; Fig. 1A) and subsequently while gamma-inducing achromatic stimuli were shown (M2). We showed achromatic gratings before (G1) and after (G2) the subjects performed stimulus-free meditation (M1). We also had protocols where participants either kept their eyes open or closed without meditation, both before (EO1, EC1) and after (EO2, EC2) the stimulus-free meditation segment (M1). Eye position was measured using a head-restraint free eye tracker (Eyelink duo; SR-Research). A fixation spot was shown for all eyes-open segments (EO1, EO2, G1, G2, M1, M2), and epochs where the eye position deviated by more than 2.5 degrees around fixation were removed offline later.

**Figure 1:**
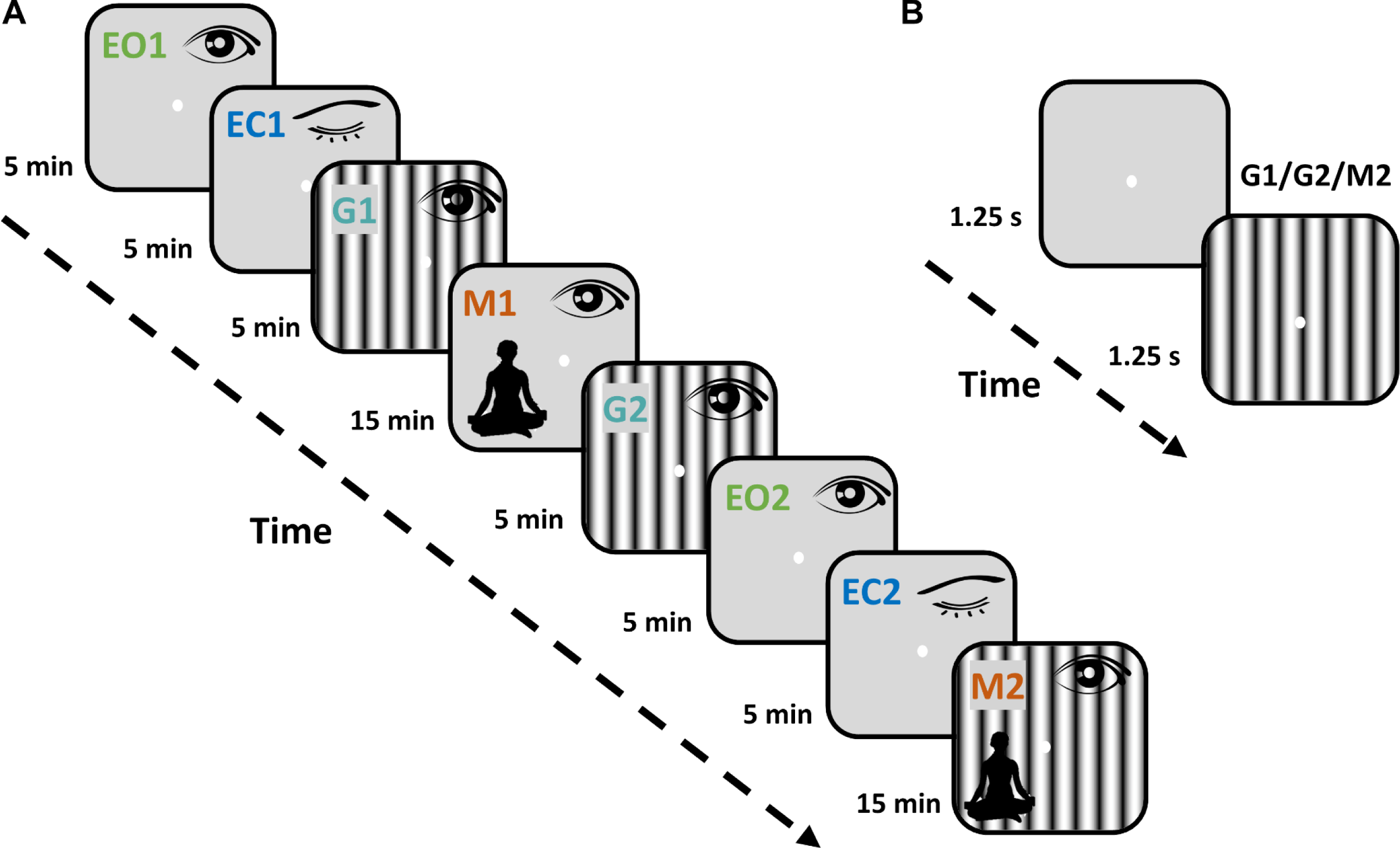
Experimental Design and the Task Paradigm. **(A)** The experiment consisted of 8 protocols, starting with eyes open (EO1) condition where subjects were required to maintain fixation on a dot on the screen. The eye position of the subjects was monitored using a restraint-free eye tracker for all protocols where eyes were open, and epochs where the eye position deviated beyond 2.5 degrees from fixation were discarded offline later. EO1 was followed by the eyes closed (EC1) condition. Gamma protocols (G1 and G2) were conducted before and after the stimulus-free meditation protocol (M1) where participants were asked to meditate while keeping their eyes open. The experiment was concluded with another set of eyes open (EO2) and eyes closed (EC2) protocols, followed by a second meditation protocol where visual stimuli were shown while participants meditated keeping their eyes open (M2). Respective session duration in minutes is indicated beside the corresponding rectangular box for every protocol. **(B)** Subjects were asked to look at a dot at the center of the screen in all protocols except EC1 and EC2. For G1, G2, and M2 protocols, gamma-inducing achromatic gratings (spatial frequency of 2 and 4; orientation of 0, 45, 90, 135) were presented for 1.25 seconds with an inter-stimulus interval of 1.25 seconds in a continuous sequence. The subjects were asked to blink or break fixation during the inter-stimulus interval if needed.

EEG data was collected from 78 subjects (38 meditators and 40 controls), out of which 71 subjects (35 meditators and 36 controls) with at least 40 (out of 64) good electrodes were chosen (see Supplementary Fig. 1 for electrode and demographic details). We further paired one control (with age within 2 years) of each meditator to get 30 matched pairs. We show results for pairwise comparison; similar results were obtained for unpaired comparison as well. We show PSDs for occipital and fronto-temporal groups (electrodes indicated in Fig. 2D) because those electrodes show strong stimulus-induced and stimulus-free meditation-induced gamma, respectively. For each protocol, we rejected subjects with less than 30 good trials and 3 good electrodes for each electrode group, so the number of matched pairs varied between 27-30 across protocols and electrode groups.

**Figure 2:**
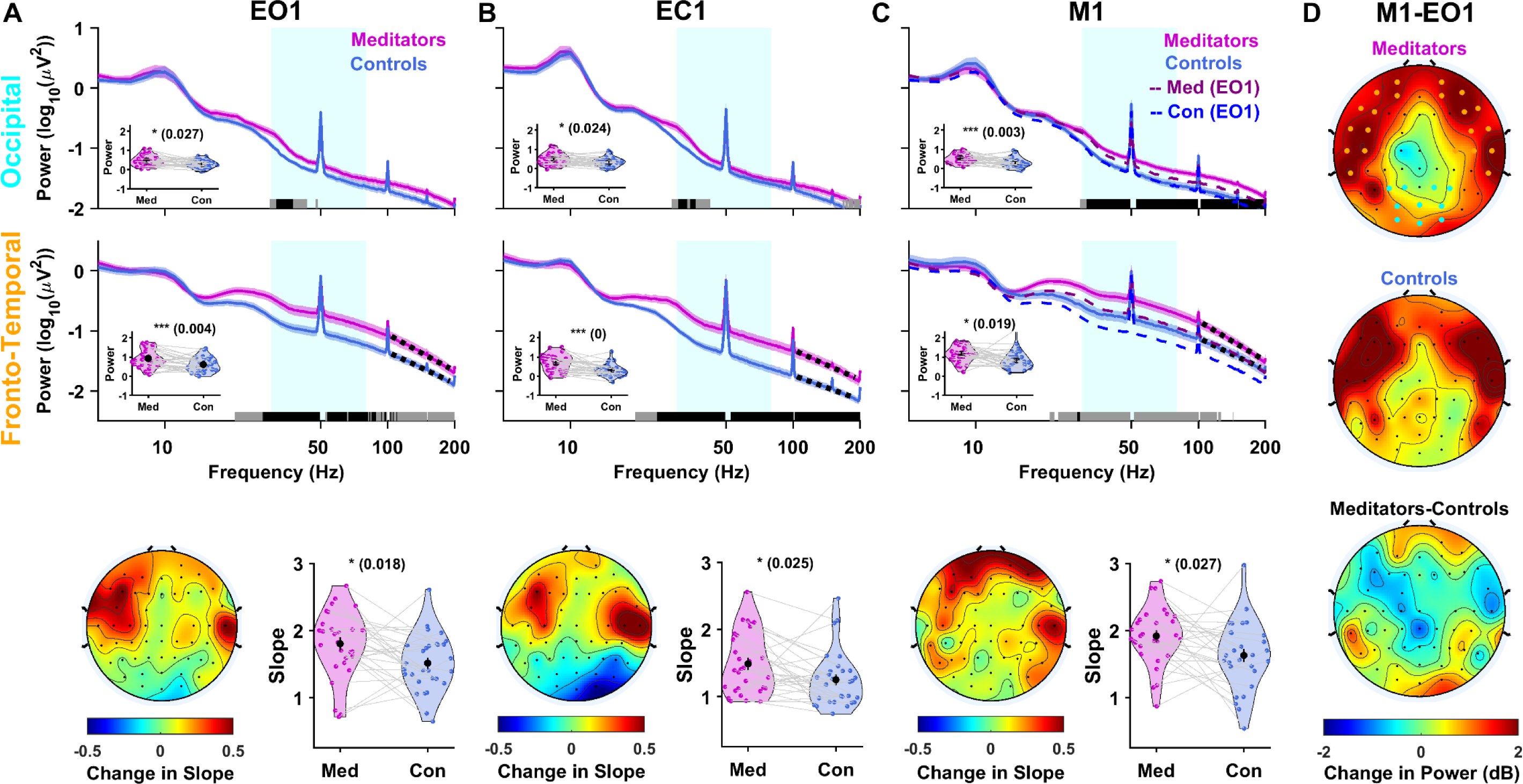
Meditators have more stimulus-free gamma during eyes-open, eyes-closed and meditation conditions. **(A)** The top two rows show the power spectral density (PSD) averaged across 29 meditators (magenta) and their matched controls (blue) for the EO1 protocol for the occipital and fronto-temporal electrode groups, respectively. Solid traces represent the mean and the shaded region around them indicates standard error of the mean (SEM) across subjects. Horizontal bars at the abscissa represent the significance of differences in mean (grey: p<0.05 and black: p<0.01, paired t-test, not corrected for multiple comparisons). The violin plots in the insets show the power for meditators and controls in the broadband gamma frequency range (30-80 Hz), shown in cyan in the main plot. Means are compared using paired t-test, and the p-values are indicated at the top, along with their significance level (*: p<0.05, **: p<0.01, ***: p<0.005). In the bottom panel, the average scalp maps (topoplots) of the 64 electrodes on the left show the change in slope between meditators and controls (meditators-controls), computed between 104-190 Hz (highlighted as black dotted lines over PSDs in the middle panel). The violin plot on the right shows the mean slope for meditators and controls in the fronto-temporal electrode group. **(B)** PSD and slope plots for the EC1 protocol when the eyes were closed. n=29 for top row and n=30 for middle and bottom rows. **(C)** PSD and slope plots when subjects meditated keeping their eyes open in the M1 protocol (n=29). Dashed lines show PSDs for the EO1 protocol for the meditators and controls (same plots as shown in A) for comparison. (**D)** Topoplots show change in gamma power (in decibels (dB)) during M1 compared to EO1 for meditators (top) and controls (middle) (n=28). The bottom plot shows the change in power in M1 relative to EO1 between meditators versus controls (difference between the top and the middle row).

### Meditators have more stimulus-free gamma during eyes-open and eyes-closed conditions

Fig. 2A shows the mean PSDs for meditators and their age and gender matched controls (n=29) for the EO1 segment for the occipital (top row) and fronto-temporal electrode groups (middle row). Even when not meditating, meditators had more power than controls over a broad frequency range above ∼20 Hz, with a shallow “bump” in the PSD around ∼30 Hz that was more prominent for meditators. This stimulus-free “broadband” gamma power (calculated between 30-80 Hz) was significantly higher in meditators for all electrodes (results for the occipital and fronto-temporal groups are shown in the insets), but more prominent in fronto-temporal regions (Supplementary Fig. 2A).

### The Slope of the power-spectral density is steeper for meditators than controls

We also compared the slope of the PSD between meditators and controls since PSD slopes become shallower with age (*25*) and steeper slopes have been associated with higher inhibition (*26*). Slopes for meditators were indeed steeper than controls over the fronto-temporal electrodes, albeit over a higher frequency range (104-190 Hz; dotted black lines in the middle row) than used previously (*25*). As shown in the scalp maps of the difference in slope between meditators and controls (Fig. 2A bottom row), this steeping of slope was mainly observed in the fronto-temporal regions.

Both the broadband change in power and the steepening of slope in fronto-temporal regions were observed when eyes were closed (EC1; Fig. 2B and Supplementary Fig. 2B), ruling out potential confounds related to myogenic activity related to open eyes.

### Meditation increases broadband gamma over fronto-temporal areas for controls, and across all areas in meditators

Stimulus-free meditation increased broadband gamma power (Fig. 2C; compare the PSDs for M1 with EO1 shown in dashed lines) in both meditators and controls (n=29). However, in occipital areas, this increase was mainly observed for meditators (Fig. 2C, top row; the separation between solid and dashed lines is only observed for meditators), while both meditators and controls showed an increase in fronto-temporal areas (Fig. 2C; middle row). To better visualize the effect of meditation across brain regions, we plotted scalp maps for the broadband gamma power (30-80Hz) during M1 relative to the power in the EO1 segment (Fig. 2D; n=28; this is obtained by subtracting the power for the two conditions on a log scale and multiplying by 10 to have units of decibels (dB)). Meditators showed an increase in both occipital and fronto-temporal areas (Fig. 2D, top row). In contrast, for controls, the increase was mainly in the fronto-temporal electrodes (Fig. 2D, middle row). This meditation-induced increase in power during M1 compared to EO1 in meditators versus controls was observed mainly in occipital and frontal regions (Fig. 2D, bottom row).

The slopes for meditators were significantly steeper than controls even during M1 (statistics shown in the bottom row of Fig. 2C). While the slopes during M1 were also slightly higher than EO1 for both groups, the difference did not reach significance (meditators: n=28 (mean±SEM) EO1: 1.80 ± 0.09, M1: 1.89 ± 0.08, p=0.12, t-test; controls: n=28, EO1: 1.50 ± 0.08, M1: 1.63±0.1, p=0.07, t-test).

### Stimulus-induced gamma is stronger for meditators even in non-meditative condition

We next examined if meditators have more stimulus-induced gamma than controls. First, we computed the PSDs for the occipital group during the spontaneous periods between the stimuli (-1 to 0 seconds; where 0 indicates stimulus onset) and found that PSDs during this epoch were similar to the EO1 condition for both G1 (n=29) and G2 (n=28) protocols (Fig. 3A; compare solid versus dashed lines), with significantly higher broadband gamma in meditators than controls as in EO1 (see insets for statistical comparison). PSDs computed during the stimulus period (0.25 to 1.25 seconds after stimulus onset; Fig. 3B) revealed a prominent narrowband gamma bump, with a peak near 25 Hz (Fig. 3B), which was more prominent in meditators versus controls for both G1 and G2 (insets). This increase, however, is unsurprising since meditators had elevated power even during spontaneous periods (Fig. 3A).

**Figure 3:**
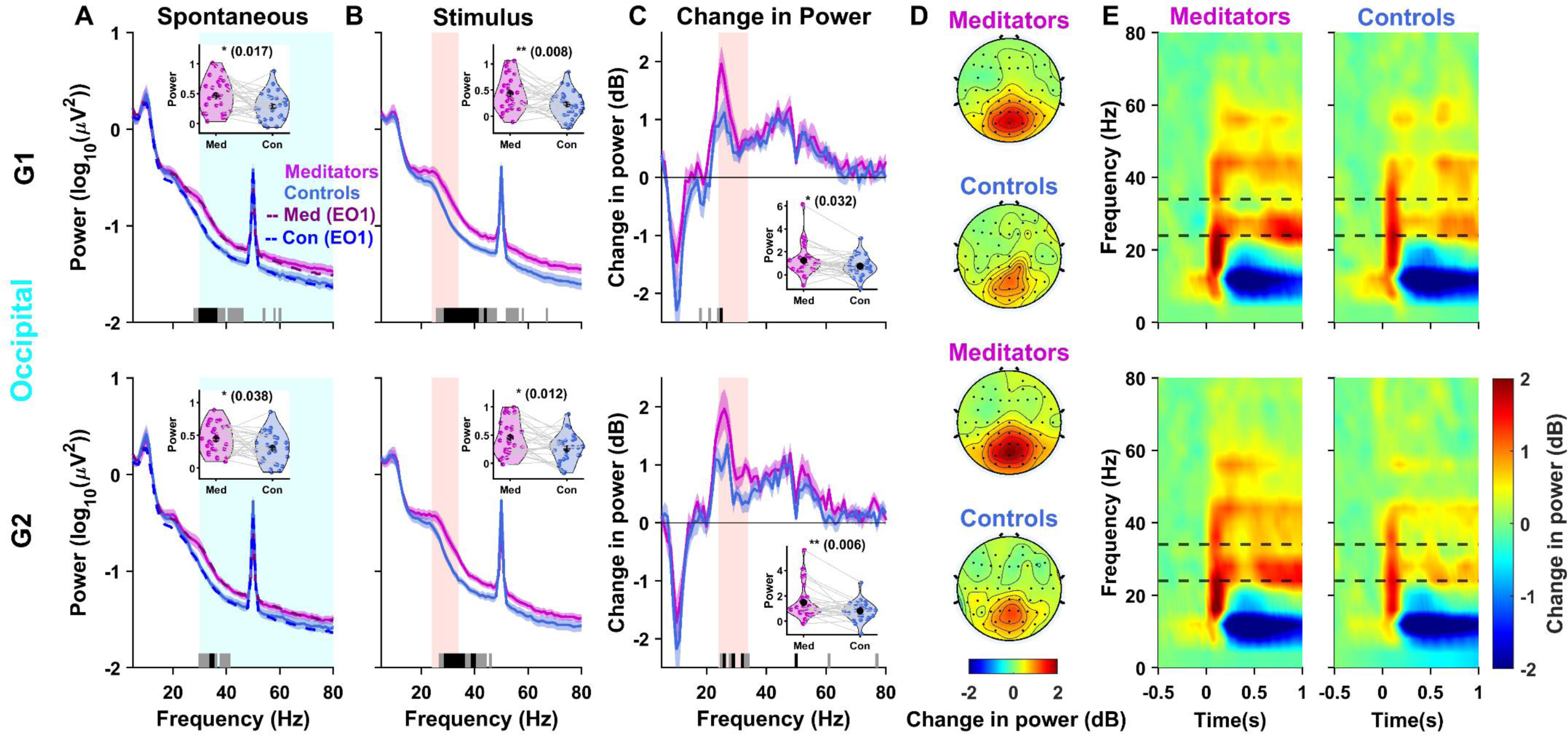
Stimulus-induced “slow” gamma is stronger for meditators. **(A)** Average PSDs across paired subjects for the occipital electrode group for the spontaneous (baseline) period (-1 to 0 seconds) before the stimulus onset for the meditators (magenta) and controls (blue) for G1 (top panel; n=29) and G2 (bottom panel; n=28) protocols. Solid traces represent the mean, and shaded regions around them indicate SEM across subjects. Dashed lines show corresponding PSDs for the EO1 protocol. Horizontal bars at the abscissa represent the significance of differences in mean (grey: p<0.05 and black: p<0.01, paired t-test). The violin plots in the insets show the broadband power difference between meditators and controls for the shaded (cyan) broadband gamma frequency range (30-80 Hz). **(B)** Average PSDs calculated for the “stimulus” period (0.25 to 1.25 seconds from the stimulus onset) for the respective protocols. The violin plots in the insets show the power difference between meditators and controls for the shaded (red) slow gamma frequency range (24-34 Hz). **(C)** Change in power in dB during the stimulus with respect to the baseline for the G1 (n=29) and G2 (n=27) protocols. **(D)** Topoplots for meditators (top) and controls (below) show the change in power during stimulus period from baseline in the slow gamma range during the respective protocols. **(E)** Time-frequency spectrogram for the meditators (left) and controls (right) for -0.5 to 1 second relative to the stimulus onset. The dashed lines show the slow gamma frequency range (24-34 Hz).

To test whether the visual stimuli induced stronger gamma, we computed the change in power during stimulus period compared to baseline (Fig. 3C). The change in power plots also revealed the fast gamma between 40-70 Hz which was not readily visible in the raw PSDs due to the line noise, as well as the suppression in alpha power around ∼10 Hz due to stimulus onset. Interestingly, while meditators had similar changes as controls in alpha and fast gamma bands, there was a significantly higher slow gamma increase in meditators than controls for both G1 and G2 protocols (Fig. 3C). Expectedly, this change in stimulus-induced slow gamma power was localized in the occipital regions (Fig. 3D). Time-frequency spectra (Fig. 3E) showed that the power in the slow gamma band increased over time, consistent with previous studies (*3*). Thus, even when the elevated spontaneous power is accounted for, meditators have stronger stimulus-induced slow gamma (but not fast gamma) than controls, even when they are not meditating.

### Stimulus-induced and stimulus-free gamma co-exist but are unrelated

Finally, we investigated the effect of meditation on stimulus-induced narrowband gamma (M2 protocol). First, we computed the PSDs during spontaneous periods. If the subjects were indeed able to meditate in spite of the continuous presentation of the stimuli, we expected these PSDs to be comparable to the PSDs obtained during M1, and further elevated compared to the spontaneous PSDs during non-meditative condition (for example, EO2 or spontaneous condition in G2). We found both conditions to be true for meditators: spontaneous M2 PSDs (solid magenta lines in Fig. 4A) were comparable to M1 PSDs (dashed magenta lines) and were elevated compared to the spontaneous G2 condition (dotted magenta lines; similar results were obtained if EO2 was used; n=29, occipital; n=30, fronto-temporal). For controls, the increase in broadband gamma power in fronto-temporal region during M1 (dashed blue line compared to dotted blue line in the bottom row) was not observed during M2 (solid blue line compared to dotted blue line). This could potentially be due to more interference experienced by controls than meditators due to the stimuli.

**Figure 4:**
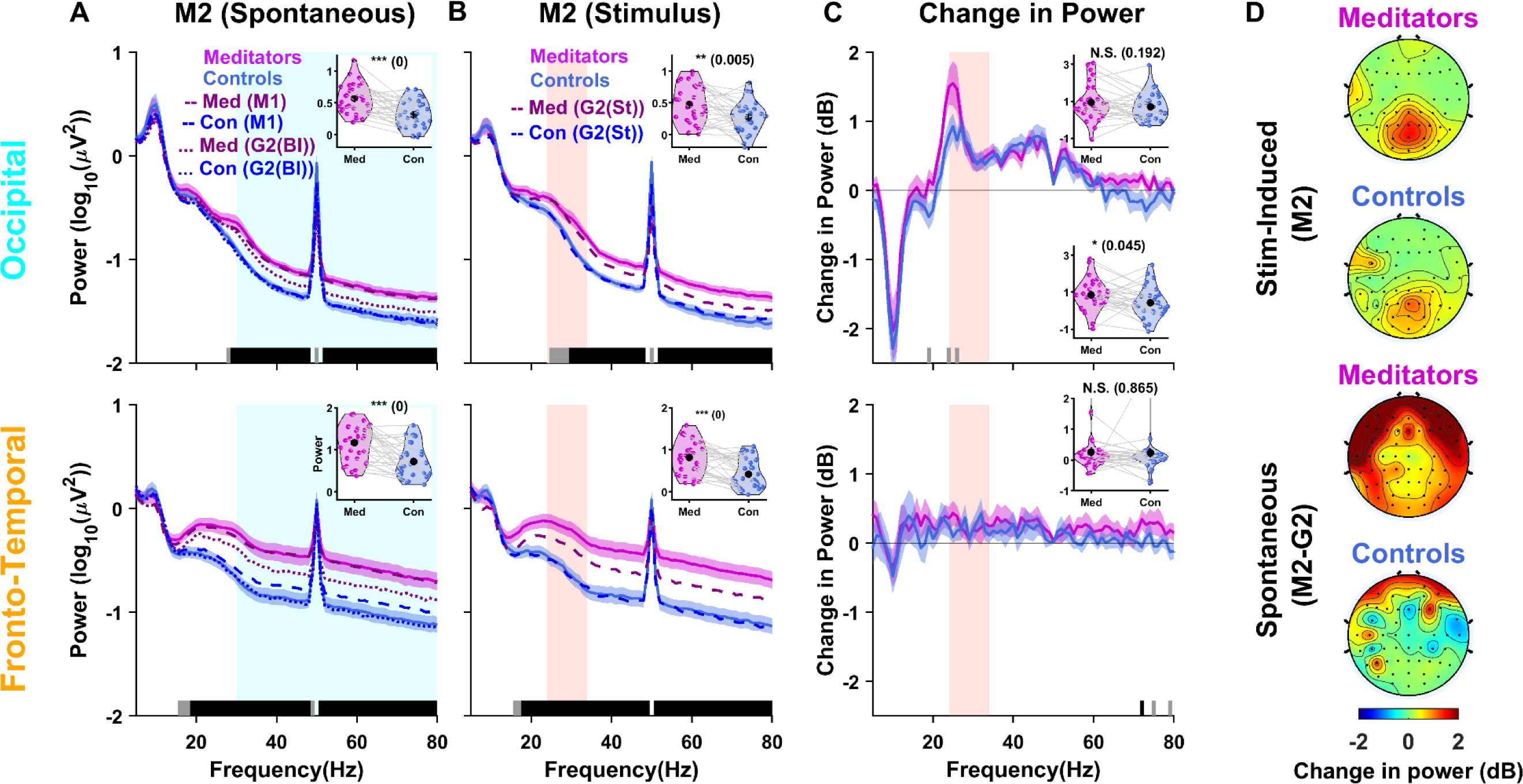
Stimulus-induced and stimulus-free gamma co-exist. **(A)** Average PSDs for the meditators (magenta) and controls (blue) for the M2 spontaneous (baseline) period for the occipital (top row; n=29) and fronto-temporal (bottom row; n=30) electrode groups (as indicated in fig. 2D top panel). Solid traces represent the mean and the shaded region around them indicates SEM across subjects. Dashed and dotted lines show corresponding PSDs for the M1 and G2 baseline (Bl) periods. Horizontal bars at the abscissa represent the significance of differences in mean (grey: p<0.05 and black: p<0.01, paired t-test). The violin plots in the insets show the power difference between meditators and controls for the shaded (cyan) broadband gamma frequency range (30-80 Hz). **(B)** PSDs for the stimulus period (0.25 to 1.25 seconds) for the M2 protocol. Dashed lines show PSDs for the G2 protocol for the stimulus (St) period. The violin plots in the insets show the power difference between meditators and controls for the shaded (red) slow gamma frequency range (24-34 Hz). **(C)** Change in power in dB during stimulus with respect to baseline for the M2 protocol. The violin plots in the insets show the power difference between meditators and controls for the shaded (red) traditional slow gamma frequency range (24-34 Hz; top) and between 20-30 Hz (bottom) **(D)** The top two panels show the topoplots for change in power during stimulus with respect to baseline during M2 (n=29), while the bottom panels show the topoplots for change in spontaneous power of M2 with respect to spontaneous power in G2 (n=27).

Presentation of the stimulus produced stimulus-induced gamma in the occipital areas (Fig. 4B), which is better observed after comparing with the spontaneous period as done previously (Fig. 4C). We again observed a higher increase in slow gamma power in meditators compared to controls in the occipital area in the slow gamma range, even though the results did not reach significance when power was computed over the slow gamma range of 24-34 Hz used earlier (Fig. 4C; top row, top inset), although it reached significance when slow-gamma was taken between 20-30 Hz (Fig. 4C; top row, bottom inset). Note that in this case, the spontaneous PSD was already elevated due to meditation and therefore stimulus-induced gamma was computed with respect to this elevated PSD. Consequently, the increase in stimulus-induced gamma was somewhat less prominent than the increase observed without meditation (compare the scalp maps shown in the top two rows of Fig. 4D, which correspond to stimulus versus baseline in M2, with the corresponding plots in G1 or G2 as shown in Fig. 3D). Also, note that the PSDs during stimulus periods of M2 were more elevated than the stimulus period of G2 (compare solid versus dashed lines in Fig. 4B) for meditators but not controls, suggesting that meditation-induced stimulus-free (broadband elevation in power) and stimulus-induced gamma (narrowband increase in slow gamma as shown in Fig. 4C) co-exist.

To find the spatial localization of meditation-induced stimulus-free gamma, we computed the change in broadband power during spontaneous periods of M2 versus G2 (difference between the solid and dotted lines in Fig. 4A) and generated scalp maps (Fig. 4D, bottom two plots). The distribution of the meditation-induced broadband gamma was similar for meditators between M2 and M1 (compare Fig. 4D third row versus Fig. 2D top row), but this increase was less prominent for controls during M2 (Fig. 4D bottom row) than M1 (Fig. 2D, second row). We tested whether subjects who had higher increase in meditation-induced gamma also had more stimulus-induced gamma. This was first tested by computing meditation-induced gamma during M1 (change in M1 versus EO1 as shown in Fig. 2D) and stimulus-induced gamma during G1/G2 (change in stimulus versus baseline as shown in Fig. 3D) and calculating the correlation. We found that the correlations were not significant between the two gamma signatures for any brain region for either meditator or control subjects. For example, Supplementary Fig. 3A shows the scatter plot for meditation-induced occipital gamma versus stimulus-induced occipital gamma (correlation (r) and p-values are indicated in the plot); similar results were obtained when we compared gamma from fronto-temporal electrodes or compared meditation-induced fronto-temporal gamma with stimulus-induced occipital gamma (data not shown). Similarly, correlations between the change in power for stimulus-induced gamma during M2 (Fig. 4D; top plots) and meditation-induced gamma in M2 (Fig. 4D, bottom two plots) were not significant (Supplementary Fig. 3B). Hence, the two gamma signatures were distinct, potentially reflecting different mechanisms of generation.

## Discussion

Our study shows that meditators have stronger stimulus-free and stimulus-induced gamma than age and gender matched control subjects. These gamma signatures can co-exist and are unrelated. Further, PSD slopes are steeper for meditators compared to controls.

Slow gamma could be generated due to the dendrite-targeting inhibitory somatostatin network, although this has been shown mainly in rodents (*14*, *15*). It is coherent over larger distances as compared to fast gamma, when recorded using both microelectrode arrays (*3*) and EEG (*21*), and therefore could be thought of a “global” gamma compared to a more local fast gamma. In comparison, the broadband increase in power, which has been observed in many meditation studies (*22–24*), has been associated with increased spiking activity in microelectrode recordings (*27*) and could reflect enhanced synchrony in macro-signals (*28*).

The slope of the PSD reflects the aperiodic “1/f” component and has been associated with many different processes such as self-organized criticality (*29*), excitation-inhibition balance (*26*, *30*), neural noise (*25*) and temporal dynamics of synaptic processes (*31*). A recent study showed steepening of slope during meditation versus non-meditative condition for meditators (*32*). We also observed a similar increase in slopes for both meditators and controls when they meditated, but the increase was not significant. The difference could be due to difference in meditation technique (closed-eye Mindfulness, Zen and Vipassana meditation versus open-eye BK Rajyoga meditation) or use of a different frequency range (2-30 Hz versus 104-190 Hz). We used a higher frequency range to avoid modulation of slopes due to oscillatory activity at lower frequencies, and because we have previously shown that the flattening of PSDs with aging can be observed at higher frequencies as well (*33*).

Overall, the increase in gamma power as well as steeper PSD slopes in meditators both provide indications of a stronger long-range inhibitory circuit, suggesting that long-term changes in the brain with meditation could protect against age and disease related neurodegeneration.

## Methods

### Subjects

We recorded electroencephalogram (EEG) data from 78 subjects who were either advanced meditators with at least 5 years of meditative practice (38 subjects; 19 female) or control subjects with no prior meditation experience (40 subjects; 17 females), with age spanning 20-65 years, from the Indian community, predominantly from Bengaluru. To recruit healthy subjects in both groups, a subjective screening was done using a health questionnaire which included information related to any chronic or current illness, any medication and menstrual cycle related information for female participants at the time of experiment data collection. Also, it was ensured by self-reporting if they did not consume alcohol, tobacco or substance of abuse in any form in the past two weeks.

The meditators were adept Rajyoga practitioners of the Brahmakumaris (BK) tradition who practiced meditation with open eyes. A questionnaire designed by the parent organization of the meditators was used to collect meditation practice data along with some factors to affirm the practice variables and dedication toward meditation practice and lifestyle. A top-down coordination from headquarters of parent organization in Mt. Abu to meditation centers in Bengaluru city was done to ensure smooth participation of the subjects. Announcements regarding the study were made by the local meditation center administrators, who also verified if the inclusion and exclusion criteria were met. Recommended subjects by local center administrators were recruited for their participation in the experiment.

Controls were recruited locally through word-of-mouth, broadcast-emails, and posters after matching their age (±2 years) and gender with meditators. For female subjects, we also tried to schedule the EEG recording such that the phase of the menstrual cycle was comparable to the meditator, since brain oscillations in the alpha/gamma band have been shown to depend on the menstrual phase (*34*). If we found a control subject first, we tried to find an appropriate meditator in a similar way. We obtained informed consents from all the participants of the study and provided monetary compensation. All procedures were approved by Institutional Human Ethics Committee of Indian Institute of Science, Bengaluru.

### Subject selection

For two meditators there was no control subject within ±2 years, and hence these two subjects were removed. For the remaining 76 subjects, we first used a fully automated pipeline (*35*) with minor modification (described below) to find bad electrodes (Supplementary Fig. 1A) and bad stimulus repeats for each subject. We rejected 5 subjects (1 meditator and 4 controls) who had more than 24 (out of 64) bad electrodes (See Artifact Rejection subsection below for more details). Therefore, the usable data consisted of 71 subjects: 35 meditators (18 male and 17 female) and 36 controls (20 male and 16 female). Finally, we paired one control subject with each meditator. In case of multiple control subjects, we chose the one who had lower difference (in order of preference) in (i) age, (ii) education level, (iii) menstrual phase (for females) and (iv) EEG recording date. For 2 male and 3 female meditators, there was no control within 2 years of age difference. Therefore, out of 35 meditators, appropriate matching control was found for 30 pairs (16 males; 14 females).

Supplementary Fig. 1B shows the demographics (age, education level, days from the first day of the last menstrual cycle for non-menopausal females) and the number of bad electrodes for all 71 subjects and the 30 matched pairs along with statistical comparison using unpaired and paired t-tests. There was significant difference between groups only for education level, since many of the control subjects were from the university campus community.

### EEG Recordings

EEG signals were recorded using 64 active electrodes (actiCAP) using BrainAmp DC EEG acquisition system (Brain Products GmbH). The electrodes were placed according to the international 10-10 system and the signals were referenced to FCz during acquisition. Raw signals were filtered online between 0.016 Hz (first-order filter) and 250 Hz (fifth-order Butterworth filter), sampled at 1000 Hz and digitized at 16-bit resolution (0.1 μV/bit). The subjects were asked to sit in a dark room in front of a gamma corrected LCD monitor (BenQ XL2411; dimensions: 20.92 × 11.77 inches; resolution: 1280 × 720 pixels; refresh rate: 100 Hz), placed at ∼58 cm from the subjects and subtended 52° × 30° of visual field for full screen gratings. Eye position was monitored using EyeLink Portable Duo head-free eye tracker (SR Research Ltd, sampled at 1000 Hz).

### Experimental Procedures

The experiment consists of 8 protocols as shown in Fig. 1. It started with passive fixation on a center dot on the screen (EO1), followed by the eyes closed condition (EC1), further followed by the passive visual fixation task (G1) where full-screen grating stimulus were presented for 1.25 seconds with an interstimulus interval of 1.25 seconds, with each protocol lasting ∼5 minutes. Stimuli were presented in an uninterrupted sequence and subjects were asked to blink (if needed) during the inter-stimulus period. Next, subjects were asked to meditate with open eyes for ∼15 minutes while keeping their gaze at the screen (M1). The next three protocols were repeats of the first three protocols (G2, EO2, and EC2). In the last protocol (M2), subjects were asked to meditate with open eyes while full screen grating stimuli were presented (like G1 and G2 protocols) for ∼15 minutes. The gamma-inducing achromatic gratings stimulus had spatial frequency of 2 or 4 cycles per degree and orientation of 0°, 45°, 90°, 135° (total 8 combinations). Even for eyes open (EO1 and EO2), eyes closed (EC1 and EC2) and stimulus-free meditation (M1) conditions, where the fixation spot remained on throughout the protocol, we segmented the data into 2.5 second “trials” involving 1.25 seconds of “spontaneous” and 1.25 seconds of “stimulus” periods to have a uniform trial structure across protocols (for protocols where no stimulus was presented, we averaged the PSDs of the two epochs). All protocols had 120 trials except for M1 and M2 which had 360 trials.

## Meditation Technique

### Meditators

The meditators practiced Rajyoga meditation (seed-stage meditation) of the BK tradition, which is practiced with open eyes. The practice employs contemplation and directed thinking in order to reach experiential states for self-development (*36*). The practice utilized in the current study was focused on shifting the awareness from the visible world to “the soul and its peaceful nature” and then connecting to the “Supreme Soul”. While practicing the seed stage, meditators started with the awareness about their body and slowly shifted their focus in the middle or behind the forehead and imagined themselves as a point of light situated in the body. This step is known as “soul consciousness” and common to all practices in BK Rajyoga tradition (*36–38*). In the final stage, meditators connect with the Supreme Soul and imagine a conversation with the Supreme Soul and establish a connection. Meditators affirm and reflect on seed like qualities of the soul (peace, power, bliss, love, purity and wisdom) and imagine receiving these qualities from the Supreme Soul which permeates the meditator’s soul. On successful connection, a feeling of completeness emerges. A feeling of detachment from the physical world emerges and meditators experience themselves in complete silence and subtleness. Seed stage symbolizes attaining original seed-like qualities seen as spiritual treasures that need to be awakened and nurtured. Meditators mentally repeat positive statements in affirmation to internalize and imbibe these qualities such as “I am a peaceful soul”, “I am a pure soul” and so on. Meditators with repeated practice can switch to seed stage in minimum time and remain in soul consciousness as a precursor.

### Controls

In order to practice relaxation beforehand, control subjects were asked to follow audio instructions. The instructions for relaxation intervention were recorded in a male voice in both English and Hindi languages, given to the recruited subject in the form of audio files and asked to practice at least once before the experimental EEG recording. Starting with body scanning, subjects were asked to focus on different body areas starting from toe of both feet and gradually moving to other parts of body and ending at the tip of the head. Apart from body scanning, they were asked to focus on the breath and sync it with the body scanning with the instruction that *“every incoming breath is energizing your body, and every outgoing breath is relaxing it”.* After this, they were asked to focus on the middle of the forehead and imagine a point of light. This procedure is similar to the technique used by BK meditators during initiation of their meditation practice. During the experimental EEG recording, they were asked to practice the same in absence of audio instructions to avoid any audio stimulation of the brain.

### Meditation experience

The meditation practice data was also collected in terms of daily meditation practice minutes in morning, evening and any other time of the day, weekly practice, and years since they started practicing it. Based on these, the total number of hours of meditative practice was calculated for all meditators except two for which this information was incomplete. Overall, the meditators had (Mean± Standard deviation (SD)) 10651 ± 7717 (n=33; Min: 2097, Max: 30576) hours of practice. We also tested whether meditation-induced or stimulus-induced gamma depended on the number of hours of meditative practice but found no significant correlation (data not shown). We note that our study was not designed to test for this, since we did not explicitly recruit meditators with varying experience. Further, stimulus-induced gamma, although consistent across multiple recordings for a given individual, varies considerably across subjects (*39*). In our data also, we found that the time-frequency plots (as shown in Fig. 3E) for G1, G2 and M2 were very similar for individual subjects (data not shown), but there was considerable variability across subjects.

## Artifact Rejection

We removed artifacts in the data using the following criteria (adapted from (*35*)). We rejected electrodes with impedance greater than 25 kΩ. Impedance of final set of electrodes was (mean ± SD) 6.19 ± 4.80 kΩ. Further, for each electrode, we detected outliers as trials with root mean square (RMS) outside 1.5-35 *μV* or with deviation from the mean signal in frequency domain by more than 6 SD, and subsequently, we discarded the electrodes with more than 30% outliers. Further, we considered trials that were deemed bad in the occipital electrode groups or in more than 10% of the other electrodes as bad, eventually obtaining a set of common bad trials for each protocol for each subject. Overall, this led us to reject (mean ± SD) 17.2 ± 6.9%, 19.2 ± 7.0%, 16.8 ± 5.5%, 23.8 ± 7.4%, 17.7 ± 6.5%, 18.5 ± 7.9%, 19.8±8.2% and 23.1±8.0% trials in EO1, EC1, G1, M1, EO2, EC2, G2 and M2, respectively. Next, we computed slopes for the power spectrum between 56 Hz and 84 Hz range for each unipolar electrode and rejected electrodes whose slopes were less than 0. We found that electrodes Fp1, FC2, FC3, FC4, FC5, FC6, FT7, TP8, C2 (indicated in Supplementary Fig. 1A) were bad in more than 35% of the subjects either in meditator or control group and hence, we excluded them for analysis in all subjects. Finally, we rejected subjects who had more than 35% bad electrodes. This led to a rejection of 5 subjects (1 meditator and 4 controls; Supplementary Fig. 1A).

### Eye-artifacts

We declared eye-blinks or change in eye position outside a 5° fixation window (i.e., ± 2.5° from the fixation spot after correcting for offset from the centre fixation spot) during −1 to 1.25 seconds from stimulus/marker onset as fixation breaks and removed them offline. This led to a rejection of a total of (mean ± SD) 31.9 ± 21.1%, 22.2 ± 19.9%, 39.0 ± 27.3%, 25.7 ± 21.1%, 33.0 ± 23.1% *and* 26.2 ± 20.0% trials in EO1, G1, M1, EO2, G2 and M2, respectively. This relatively higher proportion of fixation breaks compared to our previous studies (*18*) is because of longer analysis duration (-1 to 1.25 seconds compared to -0.5 to 0.75 seconds in previous studies) and also because the stimuli were presented in a continuous manner here as opposed to a trial-wise manner with longer inter-trial interval in previous studies. We also repeated all analyses without removing bad eye trials and found that results were similar with or without the removal of bad eye trials.

For each protocol, we rejected subject with less than 30 good trials and 3 good electrodes in each electrode group and for each epoch (stimulus or baseline). This led to a rejection of 1 subject in EO1, EC1, G1, M1, M2 and 2 subjects in G2 for the occipital electrode group, and 1 subject in EO1, G1, M1 and 2 subjects in G2 for the fronto-temporal electrode groups. Therefore, we had 30 pairs for EC1 fronto-temporal group and 29 paired subjects for all remaining comparisons except G2 for which we had 28 pairs for the analysis using both the electrode groups. For comparison across protocols (say M1 versus EO1), we selected subject pairs which were good for both protocols, leading to further reduction in the number of pairs for some comparisons (number of matched pairs varied between 27 to 30 for all comparisons).

## EEG Data Analysis

All the data analyses were done using custom codes written in MATLAB (MathWorks. Inc; RRID: SCR_001622). The analyses were performed using unipolar reference scheme. We chose two groups of electrodes, (i) fronto-temporal: a combination of frontal and temporal electrodes (Fp1, Fp2, AF3, AF4, AF7, AF8, F5, F6, F7, F8, FC5, FC6, FT7, FT8, T7, T8, C3, C4, C5, C6, TP7, TP8) to depict the effect of meditation and (ii) occipital: a combination of parieto-occipital and occipital electrodes (P3, P1, P2, PO3, POz, PO4, O1, Oz and O2) for which strong stimulus-induced gamma was observed. Scalp maps were generated using the topoplot function of EEGLAB toolbox ((*40*), RRID:SCR_007292) with standard *Acticap 64* unipolar montage of the channels.

### Power Analysis

Power spectral density (PSD) and the time-frequency power spectrogram were obtained using multi-taper method with a single taper using the Chronux Toolbox ((*41*), RRID:SCR_005547) for individual trials and then averaged across the trials for each electrode. For G1, G2 and M2, we chose “baseline” or “spontaneous” period between -1 and 0 seconds of stimulus onset, while stimulus period between 0.25 and 1.25 seconds to avoid stimulus-onset related transients, yielding a frequency resolution of 1 Hz for the PSDs. For uniformity, for other protocols where there was no stimulus (EO1, EC1, M1, EO2 and EC2), we put imaginary stimulus markers at 2.5 second intervals, chose baseline and (imaginary) stimulus periods as before, and then averaged the PSDs across baseline and stimulus. The spectrograms were obtained using a moving window of size 0.25 seconds and a step size of 0.025 seconds, thus yielding a frequency resolution of 4Hz.

We have previously shown that full-screen gratings induce two distinct gamma oscillations, which we termed slow and fast gamma (*3*). Although the peak gamma frequencies vary across individuals and also slow down with age for both gamma bands (*18*), we chose fixed frequency bands for analysis to minimize free parameters in power calculation. Since this dataset consists of younger subjects compared to our previous dataset, we chose a slightly higher frequency of 24-34 Hz for stimulus induced slow gamma (in previous studies we had subjects between 50-88 years and used 20-34 Hz (*18*, *19*)). The stimulus-free gamma range was chosen between 30-80 Hz. We calculated the change in power in the chosen frequency ranges as

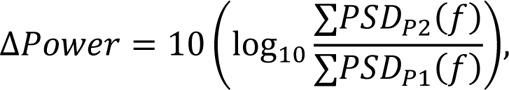

where P2 and P1 are the epochs or protocols.

### Slope analysis

We computed the slope of the 1/f aperiodic component of the spectral distribution for frequency range 104-190 Hz using the Matlab wrapper for Fitting Oscillations and One Over f (FOOOF) toolbox (*42*). The settings chosen for FOOOF model parameters were peak width limits: [4 8]; maximum number of peaks: 5; minimum peak height: 0.2; peak threshold: 2.0; and aperiodic mode: ‘fixed’, as used previously (*33*). We discarded slopes less than 0.1, which occurred due to poor fitting.

### Statistical data analysis

Since our data was matched and paired across subjects in the two groups, we compared means of the PSDs, change in power and slopes between the groups using paired t-test. Also, we verified the results by comparing the full dataset of 35 meditators and 36 controls using unpaired t-test. To find the correlation between meditation-induced gamma and stimulus-induced gamma (Supplementary Fig. 3), we calculated Pearson’s correlation coefficient (r) and p-value using the ‘corrcoef’ function in Matlab.

### Data and Code Availability

All spectral analyses were performed using Chronux toolbox (version 2.10), available at http://chronux.org. Slopes were obtained using Matlab wrapper for FOOOF (<u>github.com/fooof-tools/fooof_mat</u>). Codes to view the analysed power data is available on GitHub at <u>github.com/supratimray/ProjectDhyaanBK1Programs</u>.

## Acknowledgments

We thank the Brahmakumaris organization for their help in subject recruitment. We thank Prof. AG Ramakrishnan, Dr. Pradeep Kumar G and Adarsh A for initial discussions and help in the project.

## Funding

This work was supported by DBT/Wellcome Trust India Alliance (Senior fellowship IA/S/18/2/504003) to SR, Axis Bank PhD fellowship to AB and Prime Minister’s Research Fellows grant to SA.

## Author contributions

Conceptualization: AB, KS, SR.

Methodology: AB (power analysis), SA (slope analysis), KS (questionnaires), SR (power analysis)

Experimentation: AB (EEG), SA (EEG), KS (Subject Recruitment) Supervision: SR

Writing: AB, SA, KS, SR

## Competing interests

The authors declare no competing financial interests.

## Data and Code availability

Codes to view the analyzed power data is available on GitHub at https://www.github.com/supratimray/ProjectDhyaanBK1Programs.

## Supplementary Figure Captions

**Supplementary Figure 1.**
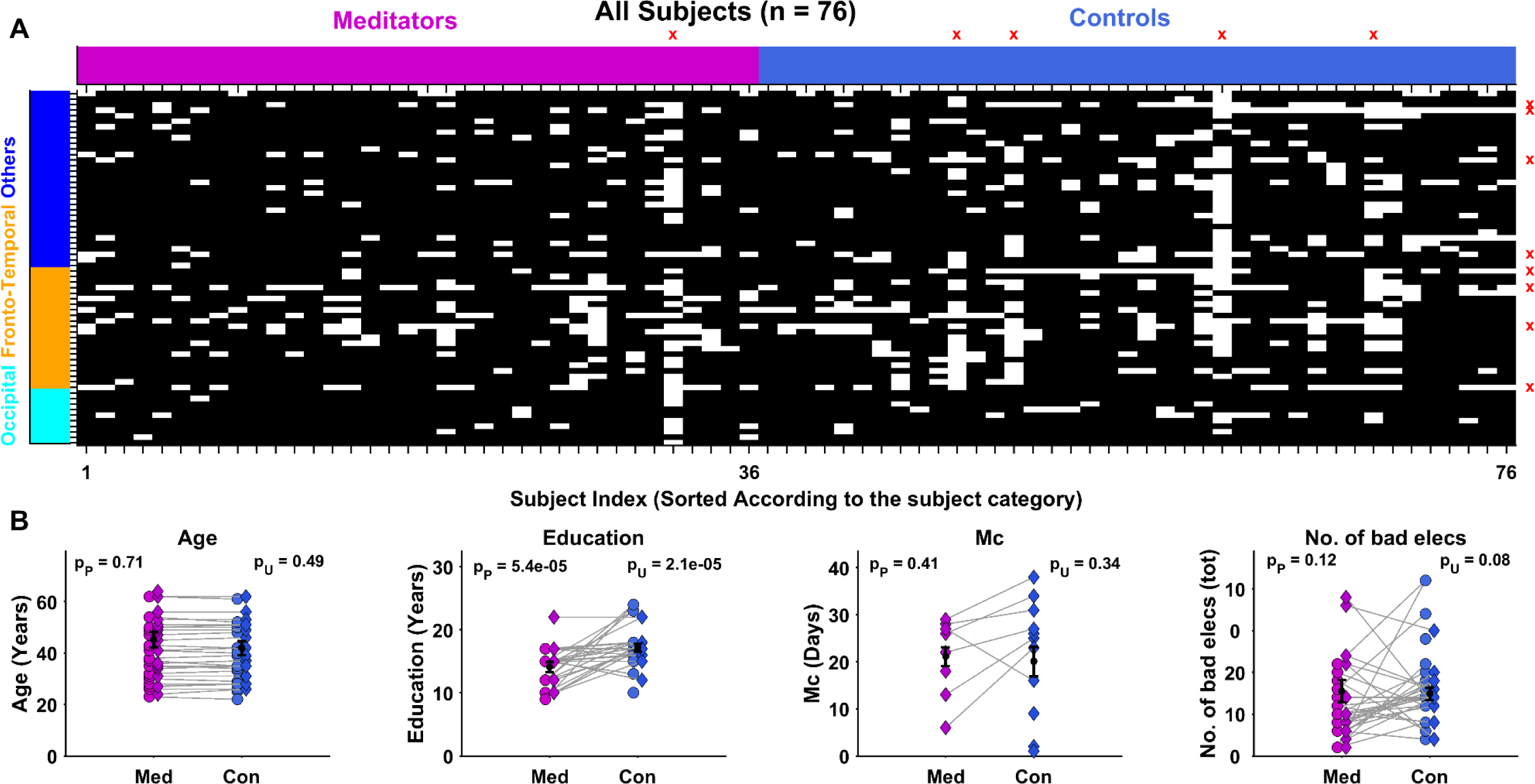
Bad Electrodes and demographic distribution of Subjects. **(A)** Heat map showing the status of 64 electrodes across all 76 subjects. The black and white colors indicate good and bad electrodes, respectively. Subjects are shown on the x-axis after grouping them as meditators or controls. The subjects with more than 40 bad electrodes are marked as red ‘x’ on the top abscissa and deemed bad (1 meditator and 4 controls). The electrodes are shown on the y-axis after arranging them in the order of occipital, fronto-temporal and remaining (“others”) electrode groups. The electrodes that were bad in more than 35% of the subjects in either meditator or control group are marked as red ‘x’ on the left ordinate and were rejected (8 bad electrodes). (B) The comparison of demographic details: age, education level, days from the first day of the last menstrual cycle for non-menopausal females and the number of bad electrodes between meditators and controls for all 71 good subjects and 30 matched pairs. The circles and diamonds represent male and female subjects, respectively. The pairs are indicated by grey lines joining the paired subjects. Means are compared using both paired and unpaired t-test, for which the p values are indicated on the top as p_P_ and p_U_ respectively.

**Supplementary Figure 2.**
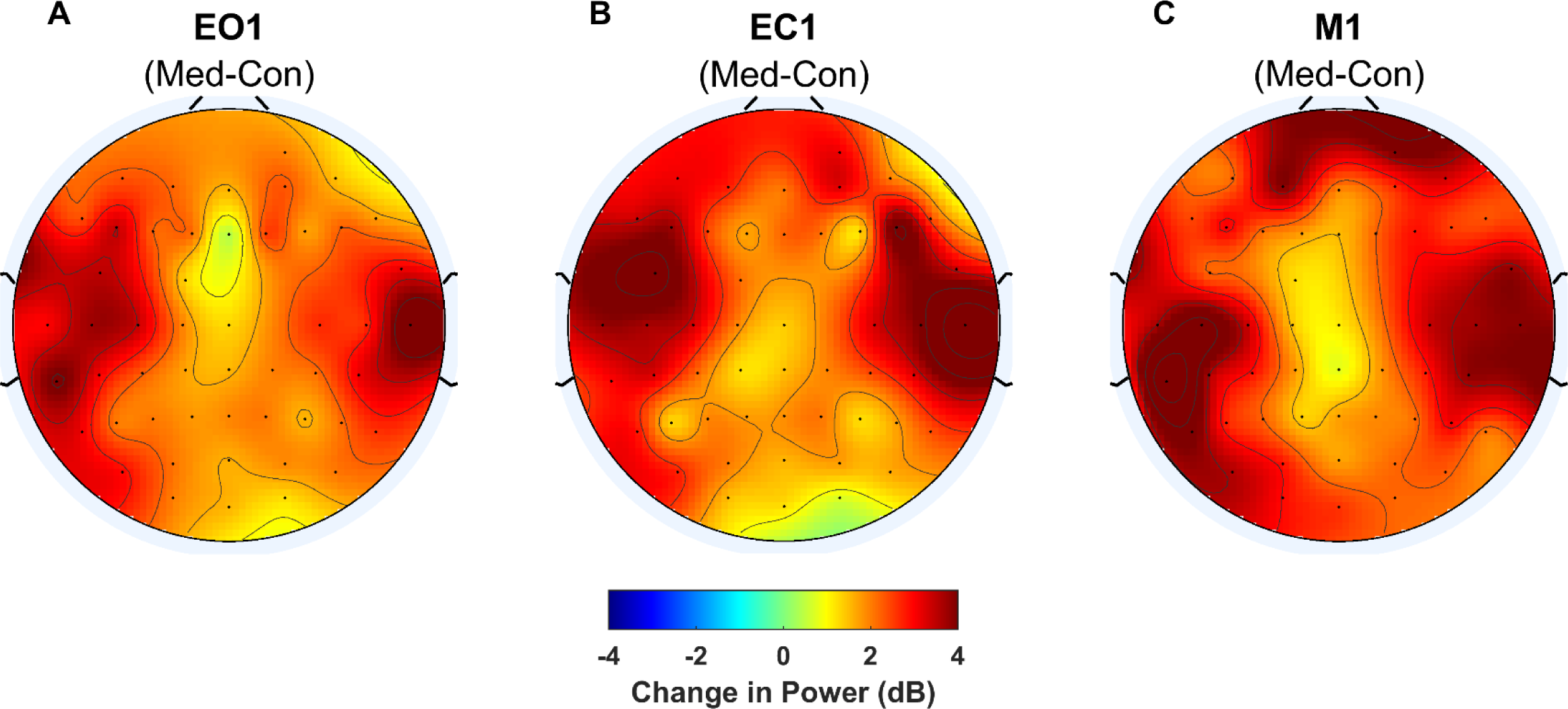
Scalp maps showing the change in power (in decibels) in the broadband gamma frequency range (30-80 Hz) for meditators with respect to controls in **(A)** EO1, **(B)** EC1 and **(C)** M1. Meditators had more broadband gamma power than controls in all brain regions.

**Supplementary Figure 3.**
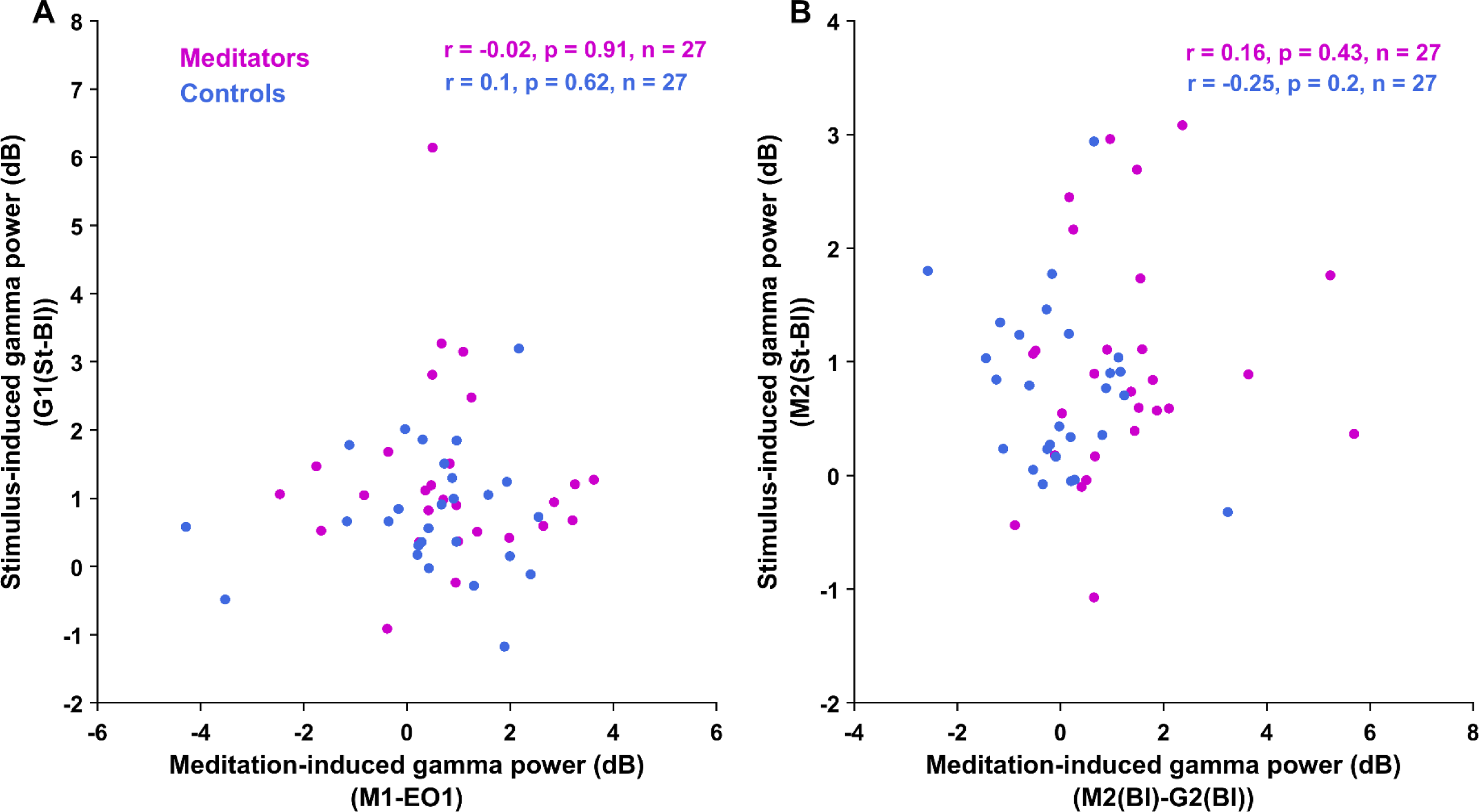
Scatter plots showing the comparison of. **(A)** meditation-induced gamma power (30-80 Hz) during M1 compared to EO1 versus stimulus-induced gamma power (24-34 Hz) during G1 stimulus compared to G1 spontaneous period **(B)** meditation-induced broadband gamma power (30-80 Hz) during M2 spontaneous compared to G2 spontaneous period vs stimulus-induced gamma power (24-34 Hz) during M2 stimulus compared to M2 spontaneous periods, for meditators (pink dots) and controls (blue dots) for the occipital electrode group. The corresponding parameters for the correlation analysis are shown on the top of each subplot for both the subject groups.

## Notes

https://www.github.com/supratimray/ProjectDhyaanBK1Programs

### Competing Interest Statement

The authors have declared no competing interest.

